# Light sheet autofluorescence lifetime imaging with a single photon avalanche diode array

**DOI:** 10.1101/2023.02.01.526695

**Authors:** Kayvan Samimi, Danielle E. Desa, Wei Lin, Kurt Weiss, Joe Li, Jan Huisken, Veronika Miskolci, Anna Huttenlocher, Jenu V. Chacko, Andreas Velten, Jeremy D. Rogers, Kevin W. Eliceiri, Melissa C. Skala

## Abstract

Single photon avalanche diode (SPAD) array sensors can increase the imaging speed for fluorescence lifetime imaging microscopy (FLIM) by transitioning from laser scanning to widefield geometries. While a SPAD camera in epi-fluorescence geometry enables widefield FLIM of fluorescently labeled samples, label-free imaging of single-cell autofluorescence is not feasible in an epi-fluorescence geometry because background fluorescence from out-of-focus features masks weak cell autofluorescence and biases lifetime measurements. Here, we address this problem by integrating the SPAD camera in a light sheet illumination geometry to achieve optical sectioning and limit out-of-focus contributions, enabling fast label-free FLIM of single-cell NAD(P)H autofluorescence. The feasibility of this NAD(P)H light sheet FLIM system was confirmed with time-course imaging of metabolic perturbations in pancreas cancer cells with 10 s integration times, and *in vivo* NAD(P)H light sheet FLIM was demonstrated with live neutrophil imaging in a zebrafish tail wound, also with 10 s integration times. Finally, the theoretical and practical imaging speeds for NAD(P)H FLIM were compared across laser scanning and light sheet geometries, indicating a 30X to 6X frame rate advantage for the light sheet compared to the laser scanning geometry. This light sheet system provides faster frame rates for 3D NAD(P)H FLIM for live cell imaging applications such as monitoring single cell metabolism and immune cell migration throughout an entire living organism.

## Introduction

Cell migration and single cell metabolism are popular areas of investigation across immunology, cancer research, and developmental biology.^1^ However, a significant bottleneck exists in 3D imaging of metabolic features in moving cells throughout a live model organism such as the zebrafish. Current methods require sample destruction (*e*.*g*., mass spectrometry, flow cytometry, histology) that removes 3D context and prevents time-course studies that follow the fate of the same cell, or fluorescent reporters that require sample manipulation.^2^ Additionally, fluorescent reporters often provide a binary on/off indicator of expression rather than a continuous variable of dynamic cell state, even though most cells are known to function on a spectrum of activity.

The reduced form of nicotinamide adenine dinucleotide (phosphate) [NAD(P)H] is a naturally fluorescent metabolic co-factor involved in hundreds of reactions within the cell (NADH and NADPH are optically indistinguishable and their fluorescence is collectively referred to as NAD(P)H).^2–13^ The fluorescence lifetime of NAD(P)H is distinct in the free (short lifetime) and protein-bound (long lifetime) conformations, so fluorescence lifetime imaging microscopy (FLIM) of NAD(P)H provides information on protein-binding activities, preferred protein-binding partners, and other environmental factors like pH and oxygen at a single cell level.^2,10,12,14,15^ FLIM of NAD(P)H is advantageous for imaging samples that are difficult to label with fluorescent reporters, for *in vivo* imaging of preclinical models, and for visualizing fast biological processes. For example, NAD(P)H FLIM can probe the temporal and spatial regulation of macrophage metabolism during tissue damage and repair in live zebrafish.^11^ However, NAD(P)H FLIM typically relies on two-photon laser scanning microscopy, which traditionally suffers from slow frame rates and limited spatial views. There is a significant opportunity to develop new NAD(P)H FLIM imaging hardware to better understand 3D cellular dynamics *in vitro* and *in vivo*.

Single-photon avalanche diodes (SPAD) are solid-state photodetectors with high photon counting and time-resolving capabilities.^16^ SPADs are semiconductor p-n junctions reverse-biased above their breakdown voltage such that a single photon event can create an electron-hole pair and trigger a self-sustaining avalanche current. Advances in SPAD technology has led to the development of multipixel arrays with on-chip integrated timing electronics (such as time-to-digital converters) capable of distinguishing single photon arrival times.^16^ SPAD arrays have numerous applications in biophotonics, including fluorescence correlation spectroscopy,^17,18^ Raman spectroscopy,^19–21^ and optical tomography.^22–24^ Larger SPAD arrays (>32×32) have also been used extensively for FLIM, which can acquire multiple emission events in parallel and eliminate the need for a scanning microscope geometry. Application examples include real time widefield FLIM of standard fluorescence solutions and quantum dots,^25,26^ fluorescently labeled cells,^27^ and fungal spores.^28^ High frame rate FLIM with a SPAD array has been used to distinguish labeled vasculature in live mice.^29^ Other geometries are detailed in the review by Bruschini, *et al*.^16^

Although SPAD arrays can be easily integrated into widefield microscopes, these microscopes lack optical sectioning capabilities and therefore offer poor sensitivity for NAD(P)H FLIM, which targets a weak autofluorescence signal over a relatively high background. Light sheet microscopy is an attractive alternative to widefield or laser scanning microscopy because it provides 3D optical sectioning along with fast volumetric imaging.^30,31^ Light sheet microscopy, also known as selective plane illumination microscopy (SPIM), uses a light sheet perpendicular to the imaging axis to illuminate the focal plane and achieve optical sectioning.^30^ Light sheet FLIM has been performed on GFP-labeled cancer cell spheroids and *C. elegans* using multichannel plate photomultiplier tube (MCP PMT) detectors,^32^ live transgenic zebrafish using a frequency-domain two-tap CMOS FLIM camera,^33^ and using a gated optical image intensifier (GOI) and CMOS camera.^34^ However, to our knowledge NAD(P)H FLIM has not been performed with light sheet systems.

Here, we demonstrate the first use of a commercially available SPAD array for detecting NAD(P)H autofluorescence in a light sheet geometry. To improve image quality, the SPAD image from the FLIM arm of the light sheet system was upscaled with the CMOS image from the intensity arm of the light sheet system. The sensitivity of the NAD(P)H light sheet FLIM system was confirmed with metabolic perturbations to pancreas cancer cells with 10 s integration times, and the NAD(P)H light sheet FLIM system was demonstrated *in vivo* with live neutrophil imaging in a wounded zebrafish tail, also at 10 s integration times. Finally, the theoretical and practical imaging speeds for NAD(P)H FLIM are compared across laser scanning and light sheet geometries, indicating a 30X to 6X frame rate advantage for the light sheet compared to the laser scanning geometry.

## Methods

### SPAD array

A FLIMera SPAD array camera (Horiba Scientific) was used for NAD(P)H FLIM.^35^ The array consists of 192×128 pixels with dedicated TDC electronics per pixel (39 ps resolution) for time-correlated single-photon counting (TCSPC). The SPAD pixels have a pitch of 9.2 µm vertically and 18.4 µm horizontally (1.75mm x 2.35mm sensor size), a fill factor of 13%, and median dark count rate (DCR) of 35 counts per second (cps). The instrument response function (IRF) of the device has a full width at half maximum (FWHM) of 380 ps. The FLIMera was controlled using Horiba EzTime Image software, with raw photon arrival time data streamed for 10 second detection periods to HDF5 files for post-processing. This data streaming mode constrained frame rates by creating a time delay between successive frames (due to EzTime software only saving data in photon stream mode, requiring 32 bits per photon event, resulting in multi-gigabyte saved files as opposed to the equivalent 3D histogram which would require a few megabytes). Conversely, integration times per image were constrained by the fill factor of the sensor (*i*.*e*., detection efficiency).

### Multidirectional selective plane illumination microscopy (mSPIM) with SPAD array

The mSPIM system is detailed in Fig. 1,^36^ with multidirectional capabilities maintained in the intensity arm only. Illumination was achieved using two opposing light sheets. Three 10x/0.3 NA water immersion objective lenses (CFI60, Nikon Instruments, Inc.) were used for illumination and detection. The “Flamingo” T-SPIM design^37^ was modified to incorporate a QuixX 375-70PS picosecond-pulsed diode laser (Omicron-Laserage Laserprodukte GmbH) in the right illumination arm for NAD(P)H excitation. The laser produced 90 ps pulses at a 50 MHz repetition rate with an average power of 0.4 mW at the sample. The output beam had a diameter of 1 mm which was used along with a f = 100 mm cylindrical lens (Thorlabs) to underfill a 10x/0.3NA objective lens to generate a 200 µm-wide light sheet with 9 µm waist thickness to match the FLIMera sensor size (right arm, Fig. 1). A bank of continuous wave lasers (TOPTICA) was used to generate a 1.5 mm-wide light sheet (left arm, Fig. 1) for targeting other fluorescent labels for intensity imaging using a larger sCMOS sensor (13.3mm x 13.3mm) with 2048×2048 pixels, 6.5 µm pitch (Panda 4.2, PCO GmbH). Brightfield trans-illumination was achieved using a red LED and imaged using the sCMOS sensor.

**Fig. 1.**
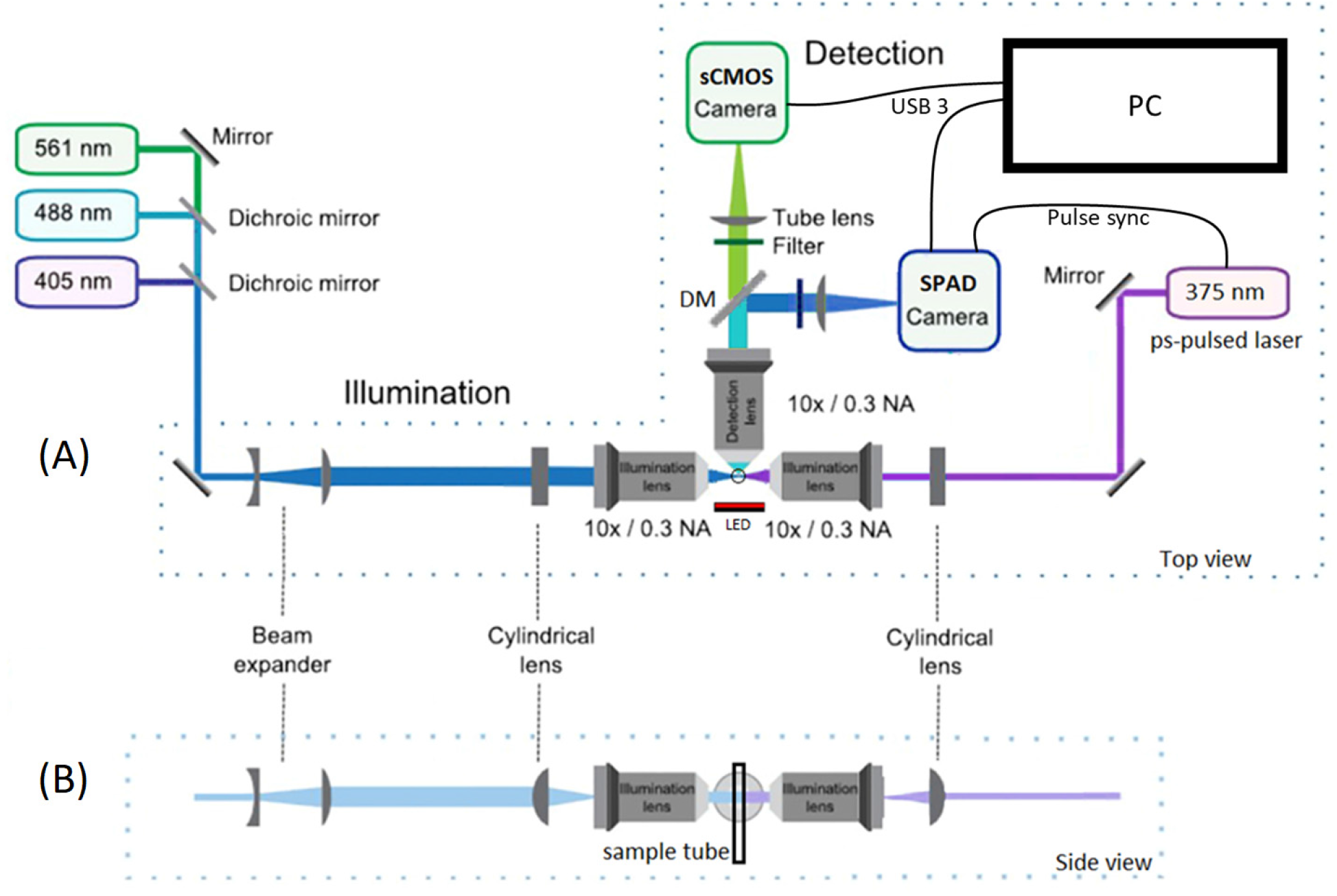
Schematic of the light sheet FLIM microscope (adapted from^36^). (A) Top view of the SPIM system. The right illumination arm uses a 375 nm ps-pulsed diode laser operated at 50 MHz repetition rate to excite NAD(P)H fluorescence via a 200 µm wide light sheet, while the time-resolved SPAD camera images the NAD(P)H autofluorescence lifetime (after a 495nm LP dichroic mirror and a 440/80nm filter) and the sCMOS camera images the NAD(P)H intensity (using a 530/55nm filter). The left illumination arm uses a bank of CW lasers to excite other fluorophores via a 1.5 mm wide light sheet, while emissions are imaged on the sCMOS camera. A red LED provides trans-illumination for bright field imaging. (B) Side view of the beams in the illumination arms and the orientation of the sample tube. DM, dichroic mirror.

NAD(P)H was excited at 375 nm (0.4 mW, 50 MHz, integration time = 10 s on SPAD array) and mCherry at 561 nm (CW laser, integration = 1 s on sCMOS camera). Emissions were split by a 495nm long-pass dichroic mirror; NAD(P)H was captured using a 440/80 nm bandpass filter on the SPAD array and a smaller NAD(P)H intensity signal was collected through a 530/55 nm bandpass filter on the sCMOS camera (integration time = 1 s). Fluorescence from mCherry and brightfield images were captured through a 650/60 nm bandpass filter on the sCMOS camera using illumination from the 561 nm CW laser light sheet or the red LED brightfield, respectively.

### Data processing

The HDF5 photon stream files were converted to 3D histograms of photon arrival times in MATLAB (MathWorks). The histograms were pre-processed to correct for the SPAD sensor artifacts^38^ *i*.*e*., variable DCR and timing skew across the sensor pixels. The DCR was estimated for each pixel from the tail of the decay (*i*.*e*., the last 2 ns of the decay) as the offset variable for the lifetime fitting algorithm. The timing skew was corrected by applying a circular shift to each pixel decay histogram by a value (up to 2.5 ns) estimated from cross-correlation maximization of each pixel instrument response decay with a reference IRF. To improve lifetime fits, a binning factor of one (*i*.*e*., 3×3 neighboring pixels) was applied to the decays. The instrument response was deconvolved from the raw decay measurements in Fourier domain.^39^ To determine NAD(P)H fluorescence lifetime parameters, the pixel decays were fit to a biexponential model 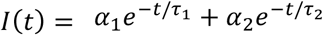. Mean fluorescence lifetime was estimated as *τ*_*m*_ = *α*_1_*τ*_1_ + *α*_2_*τ*_2_.

For visualization purposes, the NAD(P)H lifetime images from the SPAD camera were upscaled using the intensity image from the sCMOS camera acquired through the 530/55nm bandpass filter with a 1 second integration time. For upscaling, the SPAD lifetime images were interpolated to the resolution of the CMOS image (*i*.*e*., 4 times higher pixel count) and weighted by the CMOS pixel intensities. For live *in vivo* imaging of mCherry-labeled neutrophils, masks were generated from the mCherry image using a binary threshold in ImageJ,^40^ and this mask was applied to the NAD(P)H FLIM image.

### Sample preparation

PANC-1 human pancreatic cancer cells were maintained in high-glucose DMEM supplemented with 10% fetal bovine serum and 1% penicillin/streptomycin. Cells were pelleted, re-suspended in culture medium supplemented with 2% agarose and loaded into fluorinated ethylene-propylene (FEP) tubing suspended in a water bath (Fig. 1). Cells were exposed to 1 mM sodium cyanide to induce known metabolic perturbations, with images taken every 3 minutes.

### Ethics statement

Animal care and use was approved by the Institutional Animal Care and Use Committee of University of Wisconsin and strictly followed guidelines set by the federal Health Research Extension Act and the Public Health Service Policy on the Humane Care and Use of Laboratory Animal, administered by the National Institute of Health Office of Laboratory Animal Welfare.

### Zebrafish husbandry

All protocols using zebrafish in this study have been approved by the University of Wisconsin-Madison Research Animals Resource Center (protocols M005405-A02). Adult zebrafish were maintained on a 14 hr/10 hr light/dark schedule. Upon fertilization, embryos were transferred into E3 medium (5 mM NaCl, 0.17 mM KCl, 0.44 mM CaCl2, 0.33 mM MgSO4, 0.025 mM NaOH, 0.0003% Methylene Blue) and maintained at 28.5°C. Adult *casper* fish^41^ and wild-type transgenic Tg*(mpx:mCherry)* in AB background^42^ were used in this study. Tg(*mpx:mCherry*) fish were outcrossed to *casper* fish to generate transgenic fish without pigmentation. Larvae at 3 days post fertilization (dpf) were screened for mCherry expression using a ZEISS Axio Zoom.V16 fluorescence stereo zoom microscope (EMS3/SyCoP3; ZEISS; PlanNeoFluar Z 1X:0.25 FWD 56-mm lens) and raised to breeding age. Resulting adult fish were used for incross breeding. Larvae at 3 dpf were screened for *casper* larvae with mCherry expression using ZEISS Axio Zoom.V16 microscope and used for wounding and live imaging.

### Zebrafish wounding and mounting for image acquisition

3 dpf larvae were anesthetized in E3 medium containing 0.16 mg/mL Tricaine (ethyl 3-aminobenzoate; Sigma-Aldrich) and caudal fin transection was performed^11^ 30 minutes prior to imaging. Larvae were mounted in a 0.8-mm inner diameter fluorinated ethylene propylene (FEP) tube in 0.4% low-melting-point agarose with 0.16 mg/ml Tricaine and an agarose plug.

## Results

The accuracy of fluorescence lifetime recovery using the NAD(P)H light sheet FLIM system was verified by imaging standard fluorophores. Lifetimes of 2.5 ns for a saturated solution of coumarin 6 (Sigma-Aldrich) in ethanol, and 2.2 ns for 10 µm YG fluorescent microspheres (Polysciences) were recovered, which agree with reference values.^43,44^

NAD(P)H FLIM images of live PANC-1 cells were acquired using the SPAD camera and a 440/80nm filter with an integration time of 10 s to ensure sufficient photon counts (>500 photons per pixel before binning). An NAD(P)H intensity image was also acquired using the sCMOS camera and a 530/55nm filter with a 1 s integration time to upscale the SPAD FLIM images for better visualization. After the addition of 1 mM cyanide, images were taken every 3 min (Fig. 2). We observed a decrease in NAD(P)H mean fluorescence lifetime (*τ*_*m*_, Fig. 3A-D, I), an increase in the free fraction of NAD(P)H (*α*_1_, Fig. 3E-H, J), and an increase in the NAD(P)H fluorescence intensity (Fig. 3K), starting at 3 minutes post exposure and continuing over time. These changes in NAD(P)H intensity and lifetime parameters were expected,^4^ and the range of NAD(P)H *τ*_*m*_ and *α*_1_ also fall within published values for cells in culture.^45,46^ These experiments confirm the sensitivity of this system to NAD(P)H lifetimes and to physiologically relevant changes in NAD(P)H lifetimes. These studies also demonstrate that this NAD(P)H light sheet FLIM system can perform time-course imaging of single-cell autofluorescence in a fluorescent background of culture medium. Further, we found that 10 s integration time was sufficient to capture enough NAD(P)H photons for accurate lifetime fitting analysis.

**Fig. 2.**
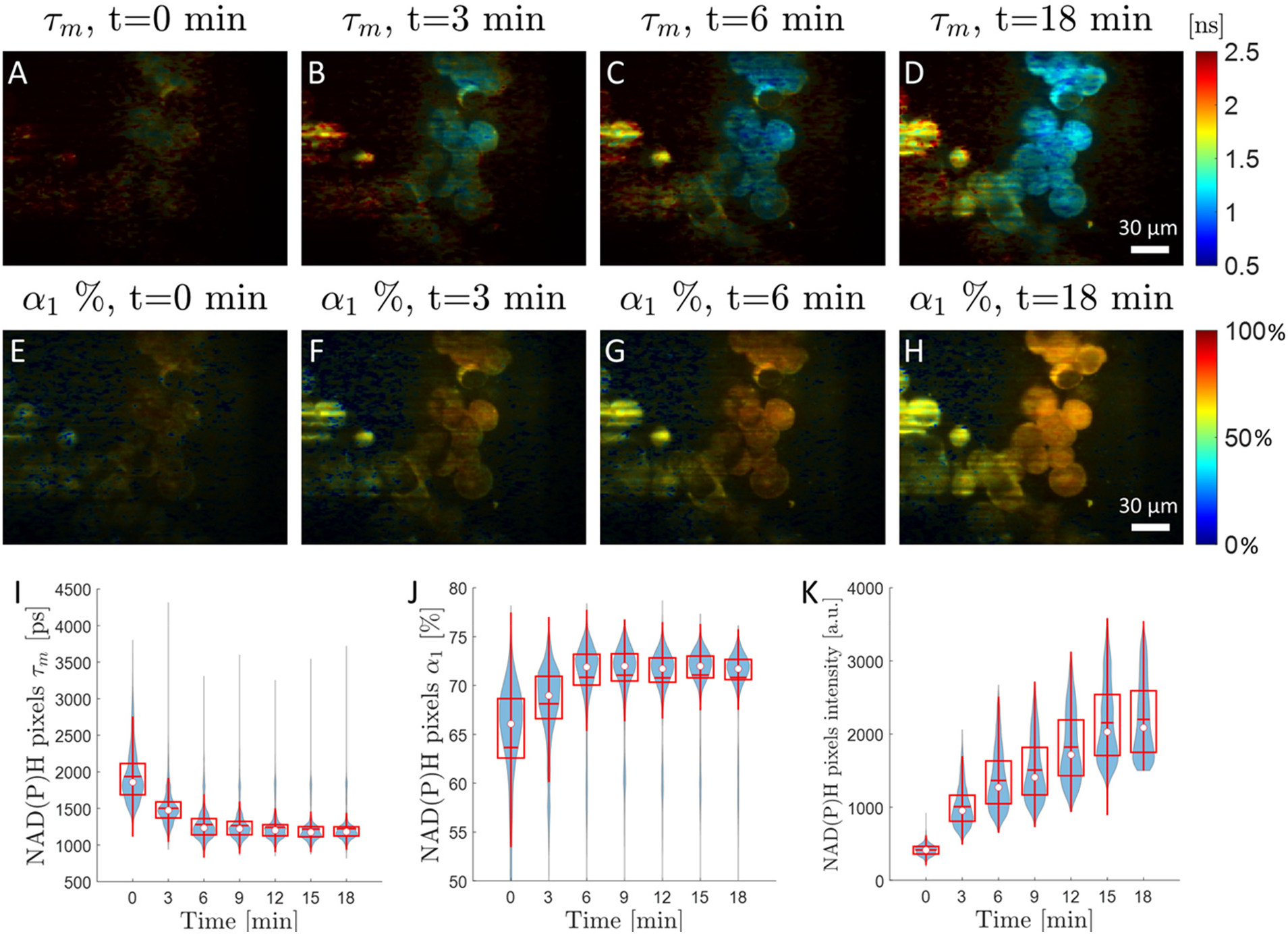
Cyanide treatment of PANC1 cells confirms the sensitivity of the light sheet FLIM system to NAD(P)H fluorescence lifetime changes. (A-D) NAD(P)H mean fluorescence lifetime (*τ*_*m*_) images of PANC1 cells at different time points after treatment with 1 mM sodium cyanide. The sCMOS NAD(P)H intensity image is used to upscale the SPAD array NAD(P)H lifetime image (E-H). Corresponding free fraction of NAD(P)H (*α*_1_,), at the same time points. (I) Boxplot of NAD(P)H *τ*_*m*_ of image pixels over time shows a rapid drop in mean lifetime within a few minutes of cyanide treatment. (J) NAD(P)H *α*_1_ of image pixels increases over time. (K) The pixel intensity of NAD(P)H fluorescence, measured by integrating the decay curve at each pixel in the SPAD camera, increases over time with cyanide exposure. White dot shows the median; red horizontal line shows the mean; box encompasses 25th to 75th percentile range; whiskers extend from the box to 1.5 times the interquartile range. Changes in NAD(P)H mean lifetime (*F*-statistic = 9.07, *p* = 0.030), free fraction (*F*-statistic = 6.88, *p* = 0.047), and mean intensity (*F*-statistic = 104, *p* = 0.0002) with time are significant according to the linear trend test.

**Fig. 3.**
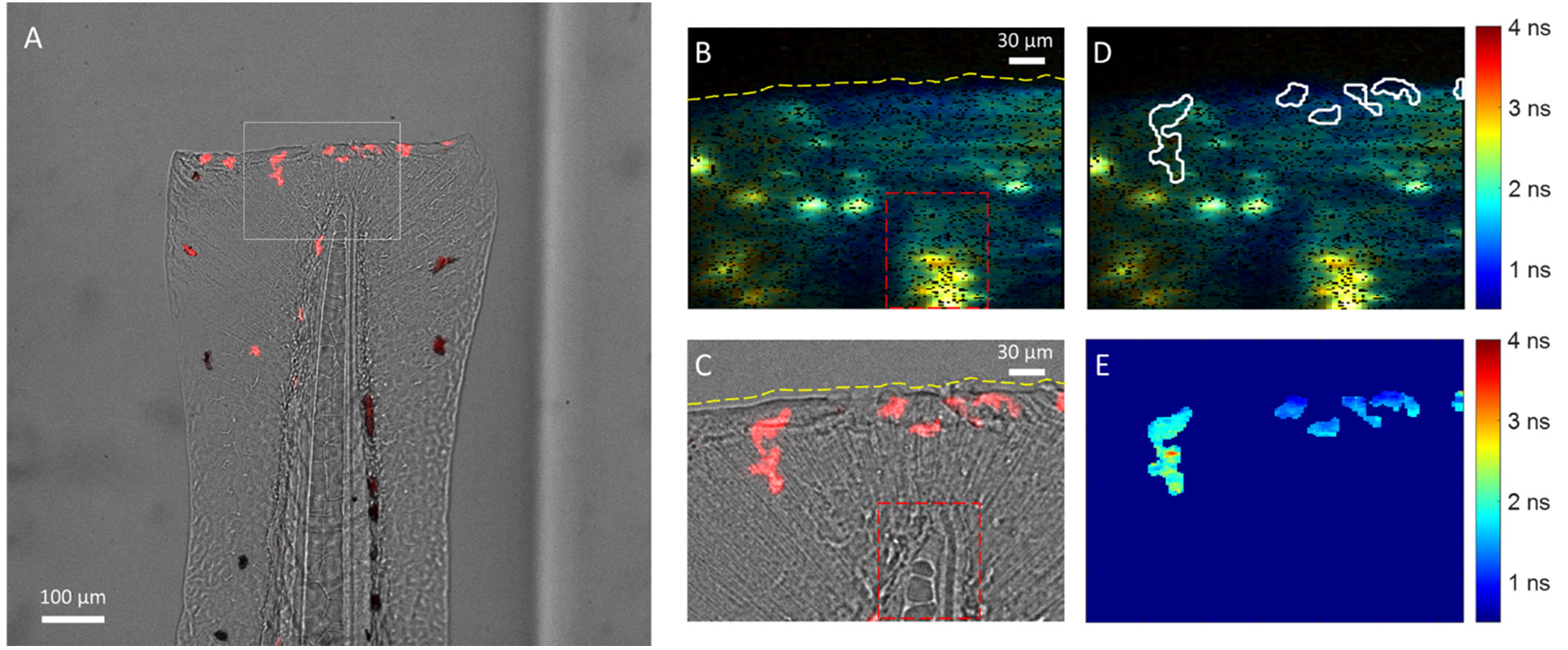
Light sheet imaging of mCherry intensity and NAD(P)H lifetimes in live neutrophils *in vivo* in a zebrafish wound model. (A) The intensity channel of the light sheet FLIM system shows the superimposed brightfield image of 3 dpf wounded zebrafish tail and fluorescence of mCherry+ neutrophils at the wound site. The white rectangle shows the corresponding field of view of the smaller SPAD array sensor. (B) NAD(P)H mean fluorescence lifetime (*τ*_*m*_) image from the SPAD of the tail (10 s integration). The transection wound (yellow dashed line) and the tip of the notochord (red dashed box) are visible in the lifetime image. (C) Corresponding field of view from the cropped brightfield image shows mCherry+ neutrophils recruited to the wound site. (D) Pixel mask of neutrophil cells from the mCherry channel applied to the NAD(P)H lifetime image. (E) Masked NAD(P)H lifetime map of the neutrophils.

We next sought to measure NAD(P)H lifetime in a dynamic, *in vivo* environment (Fig. 3). We used a zebrafish wound model to track rapidly moving neutrophils^47^, innate immune cells recruited to the injury site. We collected brightfield (25 ms exposure sCMOS), mCherry intensity (1 s exposure sCMOS), and NAD(P)H lifetime (10 s integration SPAD) from neutrophils in quick succession (every 2 minutes). NAD(P)H FLIM was acquired in a z-stack with 4 slices through the thickness of the tail fin, and all four slices show the same neutrophils. The sCMOS mCherry image was superimposed on the NAD(P)H fluorescence lifetime image to generate neutrophil masks, and lifetime information was extracted from single cells. These studies demonstrate that the NAD(P)H light sheet FLIM system is sensitive to single cells *in vivo*, and that combined intensity and FLIM images can be used to isolate quickly moving cells *in vivo* in 3D.

## Discussion and conclusion

FLIM frame rates improve for widefield detection compared to laser scanning multiphoton or confocal microscopy. Recent developments in SPAD array sensor technology enable widefield FLIM with high TCSPC temporal resolution. These new SPAD cameras can be combined with low-cost pulsed diode lasers for rapid FLIM. Table 1 shows the theoretical advantage in integration time for the SPAD array used in this work over a standard laser scanning FLIM microscope using a single TCSPC module for acquiring an image of the same size. However, the simplest implementation of widefield FLIM, *i*.*e*., an epi-fluorescence geometry, can only be applied to bright fluorescently-labeled samples due to lack of optical sectioning. In label-free applications, out-of-focus background fluorescence can overwhelm the weak autofluorescence signals from single cells. Thus, wide field autofluorescence FLIM has remained impractical.

**Table 1.** Comparison of theoretical FLIM imaging speed limit between laser scanning and widefield geometries. The saturation count rate for the SPAD camera with per-pixel TDC modules working in parallel is 30X higher than the saturation count rate of a single TCSPC module in laser scanning microscopes. The speed advantage scales with the number of array pixels.

|  | Laser scanning<br>(PMT + TAC) | Wide field<br>(SPAD array + TDC) |
| --- | --- | --- |
| Image pixel count | 192×128 = ~25000<br>sequential | 192×128 = ~25000<br>simultaneous |
| Detector dead time | ~100 ns | ~80,000 ns |
| Max pixel count rate | ~10×10<sup>6</sup> cps | ~12,000 cps |
| Max system count rate | ~10×10<sup>6</sup> cps | ~300×10<sup>6</sup> cps |
| Effective pixel count rate | ~400 cps | ~12,000 cps |

In this work, we combined multi-channel detection using a SPAD camera with light sheet excitation to minimize the effects of out-of-focus background fluorescence. In addition to providing optical sectioning, light sheet illumination is more efficient at exciting fluorophores than epi-illumination because photons traveling in the image plane have a higher probability of encountering a fluorophore. The peak irradiance of a light sheet is also many orders of magnitude lower than a diffraction-limited spot in laser scanning (*e*.*g*., confocal) microscopy, which decreases phototoxicity.

Specifically, for autofluorescence FLIM, the low quantum yield of endogenous fluorophores (*e*.*g*., NAD(P)H) is the limiting factor for increased frame rates because photodamage to cells occurs before the detector count rate saturates. As such, it is helpful to compare the light sheet SPAD system to a laser scanning system for the imaging settings used and count rates observed for NAD(P)H FLIM. This comparison is made in Table 2. For the same field of view, number of image pixels, and targeted photon count per pixel, the light sheet SPAD system in Fig. 1 requires 6X less integration time to acquire the NAD(P)H image compared to a two-photon (2P) laser scanning microscope. Frame rates of SPAD arrays relative to single-pixel laser-scanning systems scale with the number of array pixels, when acquiring the same number of image photons. SPAD arrays are based on the same CMOS fabrication process as existing cameras, so the commercial infrastructure for inexpensive and large-scale manufacture of high-resolution arrays exists.^48^ This has resulted in a rapid improvement in the capabilities of SPAD systems with improved photon detection probabilities and fill factors (*e*.*g*., using micro-lens arrays over SPAD pixels^49,50^), which can further increase FLIM frame rates or reduce excitation light dose for reduced phototoxicity over long imaging sessions.

**Table 2.** Comparison of practical FLIM imaging speed limit between a standard 2P laser scanning system and the light sheet system in Fig. 1. For the same field of view size, number of image pixels, and targeted number of emission photons per image pixel, the light sheet system required 6X less integration time per image.

|  | **2P laser scanning**<br>**(PMT + TAC)** | **1P light sheet**<br>**(SPAD array + TDC)** |
| --- | --- | --- |
| **Laser power** | 5 mW at 750nm<br>ø1µm diffraction-limited<br>Obj. 40× / 1.15NA | 0.4 mW at 375nm<br>200µm × 9µm sheet<br>Obj. 10× / 0.3NA |
| **Peak irradiance** | ~640,000 W/cm<sup>2</sup> | ~28 W/cm<sup>2</sup> |
| **FOV** | 175µm × 235µm | 175µm × 235µm |
| **Image pixel count** | 192×128 = ~25000<br>sequential | 192×128 = ~25000<br>simultaneous |
| **Total integration time** | 60 sec | 10 sec |
| **Pixel dwell time** | 2.4 millisecond | 10 sec |
| **Pixel photon count rate** | ~1.3×10<sup>6</sup> cps | ~300 cps |
| **Photons per pixel** | ~3000 | ~3000 |
| **Effective pixel count rate** | ~50 cps | ~300 cps |
| **Light dose** | ~940 J/cm<sup>2</sup> @750nm | ~280 J/cm<sup>2</sup> @375nm |

Existing light sheet FLIM microscopes perform well for samples with bright fluorophores, but access to weaker autofluorescence lifetimes is challenging due to limitations in sensitivity and temporal resolution of current detector arrays. Here, we show that it is feasible to use a commercial SPAD array in a light sheet microscope for NAD(P)H FLIM of single cells *in vitro* and *in vivo*. The integration time per image for this configuration is 6X faster than that of traditional laser scanning NAD(P)H FLIM microscopes. Faster frame rates will be achieved as SPAD arrays continue to improve in detection efficiency and array size.

These studies were limited by sensor software that saved data in photon stream mode only, which required delays between successive frames. Specifically, the FLIMera camera can only transfer data in photon streaming mode over USB 3.0, with histogram compilation and lifetime analysis on the computer CPU. The process of transferring individual photon time tags costs both USB data transfer bandwidth and analysis time, which prohibits the acquisition of successive frames at the maximum speed of the hardware. Some SPAD array manufacturers use on-chip histogramming with an FPGA to substantially reduce this data transfer and analysis bottleneck, enabling larger SPAD arrays with per-pixel TDC. Alternatively, a time gating scheme with a user-selectable number and width of time gates can be used instead of TCSPC. This approach removes per-pixel TDC electronics and simplifies the design to provide larger SPAD arrays with better fill factors. This approach also allows the user to trade temporal resolution for frame rate by reducing the number of time gates. For example, time gating was employed in SwissSPAD detectors with 512×512 pixels.^27^ Therefore, as sensor sensitivity improves, parallel improvements in data streaming and analysis will be necessary to reach the full frame-rate potential of SPAD arrays for NAD(P)H light sheet FLIM

Overall, these NAD(P)H light sheet FLIM systems will be an important new tool to study single cell metabolism and migration in 3D, including *in vivo* studies of whole model organisms.

## Abbreviations

CPS: counts per second
CW: continuous wave
DCR: dark count rate
DM: dichroic mirror
FLIM: fluorescence lifetime imaging microscopy
GOI: gated optical intensifier
IRF: instrument response function
MCP: microchannel plate
PMT: photomultiplier tube
SPIM: selective plane illumination microscopy
NAD(P)H: nicotinamide adenine dinucleotide (phosphate)
SPAD: single photon avalanche diode
(s)CMOS: (scientific) complementary metal-oxide semiconductor
TAC: time-to-amplitude converter
TCSPC: time-correlated single photon counting
TDC: time-to-digital converter

## Acknowledgments

We acknowledge funding from the Morgridge Institute for Research (MCS), Carol Skornicka Chair of Biomedical Imaging (MCS), Retina Research Foundation (RRF) Daniel M. Albert Chair (MCS), Melita F. Grunow Post-Doctoral Fellow (DED)), NIH R35 GM118027 (AH), NIH 1K99GM138699-01A1 (VM), RRF Walter H. Helmerich Professor (KWE), Beckman Center for Advanced Light-Sheet Microscopy and Data Science (KWE and JH) and NIH U54 CA268069 (KWE and MCS).

